# The impact of climate change on transmission season length: West Nile virus as a case study

**DOI:** 10.1101/2025.08.01.667982

**Authors:** R.L. Fay, C.K. Glidden, J.T. Trok, N.S. Diffenbaugh, A.T. Ciota, E.A. Mordecai

## Abstract

Climate change is accelerating the spread of temperature-sensitive vector-borne diseases such as West Nile virus (WNV), the most widespread mosquito-borne disease in the United States, with 2,400 reported cases in 2024. In New York State (NYS), where WNV first emerged in the US, mean temperatures have increased by 1.4°C since the early 1900s. Although temperature is a well-established driver of WNV transmission, its effect on transmission season length remains poorly quantified. This study examines whether WNV transmission seasons have lengthened in NYS and whether these changes are associated with increased disease risk, shifts in seasonal timing, and anthropogenic climate change. We integrated daily county-level observational data from 1999–2024, including temperature, mosquito surveillance, and human case data in a mixed-effects model, with climate model simulations spanning 1860–2024 to assess trends in transmission season length. Based on observed temperature suitability, the WNV transmission season has lengthened by an average of 19.7 days over the past 25 years and 29.6 days over the past 45 years, averaged across NYS. Longer transmission seasons are positively associated with higher WNV prevalence in both mosquito vectors and humans. While such changes could occur in the absence of global warming, climate model analyses indicate that the observed increase in season length is 7.1 times as likely under observed climate forcing since 1900 than under pre-industrial conditions.

## Introduction

Global warming since the 19^th^ century has reached 1.4°C^1^, and there has been a global average increase of 0.4°C^2^ since 1999. Climate change affects many infectious diseases, and vector-borne diseases are particularly susceptible to thermal changes because mosquitoes are ectotherms^3^. Temperature affects several key mosquito life history traits—such as survival, blood feeding, vector competence, and development—as well as pathogen replication, all of which are essential for pathogen transmission^4^. Vector-borne diseases account for more than 17% of all infectious diseases, causing more than 700,000 deaths annually^5^. Vector-borne diseases are especially responsive to anthropogenic changes, and rising temperatures can influence disease risk^6^. Although a large body of research has documented impacts of temperature on vector-borne disease transmission^6–9^, very few studies have investigated how climate warming has affected the transmission season in temperate environments^10–13^, where transmission is limited by cool temperatures in the spring and fall. As a result, seasonal epidemics that start earlier and last longer can increase the overall disease burden.

West Nile virus (WNV), a mosquito-borne arbovirus, emerged in the Western Hemisphere in 1999, with New York State (NYS) at the epicenter of the outbreak. In NYS, the average temperature has increased by nearly 1.4°C from 1901 to 2022^14,15^. Based on mathematical models parameterized using data from lab experiments, WNV transmission by *Culex pipiens*, the primary vector in NYS, is possible between 16.7°C and 33.9°C^16^. This implies that the time at which seasonal temperatures become suitable for WNV transmission governs the outer bound on the transmission season, and that these seasonal thermal suitability limits may be sensitive to climate warming.

Seasonal and decadal trends in temperature, precipitation, and relative humidity are known to influence WNV transmission across the US^7,8,16–20^. Warmer temperatures, elevated humidity, and heavy precipitation contributed to an increased rate of human WNV infection in the United States (US) during the early 2000s^21^. Drought has also been identified as a key driver of WNV transmission dynamics in the US^22^. In the Southeastern US, dry and hot spring/summer conditions, along with increased temperature and rainfall in late summer/fall, have been shown to lengthen the WNV transmission season^23^. In the Northeastern US, WNV is predicted to become more geographically widespread due to climate change^12,24^. Although the impact of temperature on WNV transmission is well described^9,16,22^, its effect on the length of the WNV transmission season and its association with human incidence have not been investigated in NYS or elsewhere. Extended virus activity could influence public health responses, including surveillance duration and resource allocation.

In this study, we examine the impact of climate change on the transmission dynamics and seasonal duration of vector-borne diseases using WNV in NYS as a case study. Using temperature, mosquito surveillance, and human case data from 1999 to 2024, spanning the entire time since WNV was introduced to the US, we assess the influence of recent temperature trends on WNV transmission season length in NYS. Specifically, we address three questions: (1) Has the WNV transmission season length increased in NYS? (2) Are longer transmission seasons associated with increased rates of WNV in mosquitoes and humans? and (3) Can these observed changes in NYS WNV season be linked to human-caused climate change? The goal of this study is to characterize how rising temperatures may affect WNV transmission season length, providing insights that could help public health officials better prepare for future transmission seasons by adjusting the timing of surveillance and vector control activities. More broadly, what do these findings suggest about how continued climate warming may reshape the duration and geographic extent of vector-borne disease transmission seasons, particularly in temperate regions where cool temperatures currently constrain transmission?

## Methods

### Environmental Data

Daily temperature data were derived from GridMET^25^, a gridded surface meteorological dataset that covers the continental US and maps surface weather variables at ∼4-km horizontal resolution. This dataset combines two sources: temporally rich data from the North American Land Data Assimilation System Phase 2 (NLDAS) and spatially rich data from Parameter-elevation Regression on Independent Slopes Model (PRISM). We downloaded this dataset from https://www.climatologylab.org/ and calculated daily, county-wide mean temperatures for the period 1980–2024 using NYS county boundaries provided by https://www.ny.gov/. Data analysis was performed in R version 4.3.2^26^, Python 3.12.9^27^, and Google Earth Engine, and visualization was conducted in R and Python.

We included elevation, El Niño-Southern Oscillation (ENSO), and the Standardized Precipitation-Evapotranspiration Index (SPEI) conditions in analyses to account for heterogenous changes in season length, anomalous temperature conditions and water availability. Mean county elevation was computed for each New York county using SRTM 1 Arc-Second Global elevation data (30m resolution^28^). ENSO conditions were characterized using the Oceanic Niño Index (ONI), retrieved from the NOAA Climate Prediction Center via the *rsoi* R package^29^. Daily gridded 3-month SPEI data is available at ∼4km horizontal resolution through the GridMET dataset. We derive the county-level SPEI values following the same approach as the GridMET temperature data.

We additionally used population to standardize case counts. Annual county-level population counts for New York State (1999–2024) were derived from the Global Human Settlement Layer gridded population product (GHS-POP, release P2023A^30^). GHS-POP provides residential population estimates at 100 m resolution for five-year epochs (1995–2025). Annual population estimates were estimated by linear interpolation between the two bracketing epoch grids on a per-pixel basis, with GHSL no-data fill values masked prior to interpolation, and county totals were then calculated by summing the pixels in each county.

When evaluating epidemiological significance of model estimates, we used population-weighted elevation to capture the elevation at which county residents are likely exposed to WNV. Population-weighted mean elevation was computed using SRTM 1 Arc-Second Global elevation data^28^ and a temporal mean GHS-POP surface across the 2000, 2005, 2010, 2015, and 2020 epochs. For each county, the population-weighted mean elevation was calculated as the sum of pixel elevation × population divided by the sum of pixel population, with both sums computed within county polygons at 100 m resolution. The code used for these analyses, as well as subsequent analyses, can be found at https://github.com/Rachellfay/WNV-Season-Length.

### Mosquito Surveillance Data

Mosquito trapping and testing data across NYS were compiled as part of the Arbovirology surveillance program, which received samples from 45 of the 62 counties, although participating counties varied by year. These data excluded Bronx, Kings, New York, Queens, and Richmond counties, which conducted their own mosquito surveillance annually. *Culex* spp. mosquitoes were collected in Centers for Disease Control (CDC) light traps by NYS county health departments and pools were submitted to the NYS Arbovirus Laboratory for processing and testing. Mosquito sampling was conducted between May and October from 2000 to 2024, with sampling intervals ranging from every 2 to 8 weeks, depending on the year and location. WNV-positive mosquito pools were identified using real-time qRT-PCR assay as previously described^31^. *Culex pipiens* and *Culex restuans* are pooled before testing because they are morphologically indistinguishable. WNV prevalence was determined using maximum likelihood estimation (MLE) based on mosquito surveillance pool sizes using an Excel Add-In (https://www.cdc.gov/westnile/resourcepages/mosqSurvSoft.html). County-level graphs were made by filtering data to only include counties where mosquito surveillance occurred for 10 or more years. Data analysis and visualization were performed in R^26^.

#### Human WNV Data

County-level human West Nile virus case data for 1999–2024 were downloaded from the CDC ArboNET Historic Data portal, which provides reported human WNV disease cases by county of residence^32^. Cases were aggregated by year. Data analysis and visualization were performed in R^26^.

### Temperature-suitable transmission season

To calculate season length, temperature data were filtered to include days when the average daily temperature was ≥16.7°C—the thermal minimum for WNV transmission estimated by previous studies^9,16^—and were within a 13-day window of another day ≥16.7°C, given what is known about historical use of lags in WNV research. This filtering removed isolated single-day temperature outliers and accounted for the mosquito development time^21,33^. Season length was measured as the number of days between the first and last days within this filtered period. Season length was calculated per year from 1980-2024. Although WNV was not detected in the USA until 1999, we included temperature data from 1980 onward to provide a longer baseline, reducing the influence of year-to-year variability and outliers on trend estimates. We did not apply an upper temperature bound because daily mean temperatures did not exceed the WNV transmission maximum of 33.9°C during the study period^9^. We conducted sensitivity analyses to test the robustness of change in season length to the thermal threshold, which included re-calculating season length at the credible intervals of the thermal minima (15-20°C) as well as population-specific thermal minima and corresponding credible intervals as defined in Fay et al. 2024^16^.

We estimated temperature-dependent change in WNV transmission season length, season start date, and season end date by fitting Bayesian hierarchical models using Integrated Nested Laplace Approximation (INLA) using the R package *INLA*^34–36^. The hierarchical Bayesian framework was selected because it allows us to simultaneously account for variation across years and locations while quantifying uncertainty in the estimated relationship between temperature and WNV transmission-season characteristics. Year was the primary explanatory variable; we tested both linear and quadratic year effects, as warming could plausibly produce non-constant rates of change over time. To account for anomalous inter-annual climate variability, we evaluated three parameterizations of the El Niño–Southern Oscillation: a continuous linear term based on mean summer Oceanic Niño Index, a nonlinear random-walk smooth (RW2) over binned ONI values, and a binary El Niño indicator (ONI ≥ 0.5). All models included the classic Besag-York-Mollié 2 spatial random effect with penalized complexity priors to account for structured (e.g., unaccounted-for climatology) and unstructured (e.g. localized urban heat islands) county-level spatial variation. We tested whether the rate of temporal change varied spatially by including year interactions with county-level mean latitude, county-level mean elevation, and county-specific random slopes (IID). We compared models using three complementary fit criteria: Deviance Information criterion (DIC)^37^. Widely Applicable Information Criterion (WAIC)^38^, and Log Conditional Predictive Ordinate (LCPO)^39^, where lower DIC and WAIC and higher LCPO indicate better predictive fit.

After selecting the best-fitting model, we derived county-specific estimates of change in season length, season start date, and season end date from the start of GridMET to present (1980-2024) and time of WNV emergence to present (1999-2024). Because the best model included a quadratic time trend interacted with county elevation, county-level change over an interval [t₁, t₂] was computed analytically from the posterior mean fixed effects (Eq. 1).

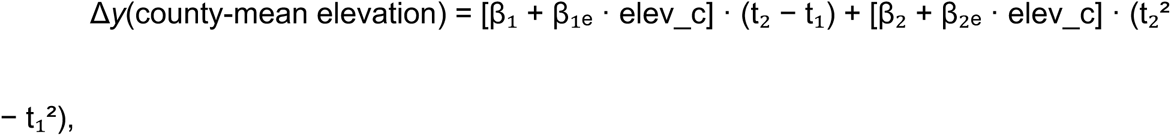

where β₁ and β₂ are the linear and quadratic year coefficients, β₁ₑ and β₂ₑ are their respective interactions with elevation, and all terms were back-transformed by multiplying by the outcome’s standard deviation. Ninety-five percent credible intervals were propagated from the 2.5th and 97.5th posterior quantiles of each fixed effect. After selecting the final model structure, we additionally assessed the sensitivity of the year effect to the temperature threshold by refitting the model using the alternative thresholds described above.

### Epidemiological Importance

To examine whether counties that experienced greater expansion of the transmission season also had a higher WNV burden, we fit two generalized additive models (GAMs) using REML estimation (R package *mgcv*): (1) total human case counts from 1999-2024 and (2) average WNV prevalence in mosquitoes from 1999-2024. Total cumulative cases per county were modeled using a negative binomial family with a log-population offset, with the 1999–2024 predicted change in season length as a penalized thin-plate regression spline (basis dimension k = 3). Mean WNV prevalence in mosquito pools per county was modeled separately using a quasibinomial family with a logit link and the same spline specification; a quasibinomial family was used to accommodate bounded proportional outcomes with overdispersion. Both models included mean summer (June–August) water availability, characterized by the summer (June-August) 90-day Standardized Precipitation-Evapotranspiration Index downloaded from GridMET and averaged over 1999–2024, as a linear covariate. Elevation was population weighted for this step of the analysis as that represents the elevation at which people are exposed.

Given the ecological-epidemiology of WNV, we would expect spatial auto-correlation in both human cases and mosquito prevalence. However, given the small analytic samples (incidence model: N = 62; prevalence model: N = 45), we tested for correlated residual spatial structure post-hoc rather than incorporate a spatial random effect, which risked overfitting at this sample size. Accordingly, Moran’s I was computed on GAM residuals using a row-standardized 5-nearest-neighbor spatial weights matrix constructed from county centroids to assess whether the fixed effects (temperature-dependent season length change and mean summer water availability) adequately accounted for the spatial patterning in each outcome. Further, given the small analytic sample sizes and the potential for individual counties to exert disproportionate influence on a spline fit we conducted a leave-one-out (LOO) sensitivity analysis to evaluate the stability of the GAM estimates. In this procedure, each county was sequentially omitted, and the model was refit on the remaining data. County-level influence was quantified by changes in deviance explained, shape of the spline, and season length significance level across leave-one-out iterations, providing a direct assessment of whether any single county was driving the estimated relationship.

### Climate model simulations

To investigate the influence of anthropogenic climate forcing on the duration of the WNV transmission season, we analyzed climate model simulations from Phase 6 of the Coupled Model Intercomparison Project (CMIP6), the most recent complete CMIP iteration. In order to weight each climate model equally, we analyzed 1 simulation per climate model (i.e., 37 simulations of the 1850-2025 period from 37 different climate models). For CMIP6, the simulations with actual historical forcing were only run through the year 2014, after which the simulations were forced by various future scenarios. Therefore, the climate model simulations we analyzed used historical climate forcing (including historical anthropogenic emissions) from 1850-2014 and, following the IPCC^41^, projected future emissions from the shared socioeconomic pathway SSP2-4.5 scenario (which is a moderate forcing scenario^41^) for the 2015-2025 period.

Since WNV transmission rates are highly dependent on absolute temperature thresholds (e.g., with 16.7°C representing the thermal minimum for WNV transmission^16^), biases in daily temperature between the GridMET dataset and the climate model simulations could substantially impact our results^42^. Therefore, we first bias-corrected and downscaled the climate model simulations to the GridMET 4km grid using the bias-correction and spatial disaggregation (BCSD) method^43^. For each climate model simulation, we used detrended quantile mapping^44^ to correct each daily grid cell temperature value to the corresponding quantile in the GridMET dataset (bilinearly interpolated to match the climate model grid). To preserve the long-term temperature trends in the climate model simulations, the long-term trends were removed (using a 30-day sliding window) from each climate model simulation and added back after the quantile mapping procedure was complete. Finally, we spatially disaggregated the climate model simulations to the 4km GridMET grid by first removing the bias-corrected climatology, then bilinearly interpolating the daily residual fields from the native climate model grid to the 4km GridMET grid, and then adding the 4km GridMET climatology back in, yielding the final bias-corrected and downscaled daily temperature data for each climate model simulation.

For each of the 37 bias-corrected and downscaled climate model simulations, we calculated a time series of daily mean temperature for each county in NYS using the county shapefiles provided by https://www.ny.gov/. We then calculated the WNV season characteristics (season length, start date, and end date) for both the pre-industrial period (1860-1904) and the recent period (1980-2024) using the same methodology as for the GridMET dataset (described in section 1 of Methods). To estimate the change in each season characteristic over a 45-year period, we fit a nearly identical INLA model to each dataset (one per GCM × time period combination), including a non-linear effect of year, an interaction between year and elevation, and a county-level spatial random effect. We then used posterior samples of the fitted year effects to calculate the change in days between the first and last year of each period, evaluated at mean county elevation given the year × elevation interaction; this yielded a posterior mean and 95% credible interval for each GCM × period combination. We did not include the ENSO effect from these models, however, we also fit an ENSO-free version of the INLA model to the observed GridMET data (1980-2024) to enable direct comparison between the climate model ensemble and the observed climate.

To quantify whether anthropogenic climate change altered the probability of the observed changes in WNV season characteristics, we compared the observed 45-year trends with the distribution of trends simulated by the climate models under preindustrial and modern climate conditions. For each season characteristic (season length, first day, and last day of the transmission season), the observed trend was defined as the 45-year change estimated from the ENSO-free INLA model fit to GridMET data for 1980–2024. The corresponding modeled trends were the posterior mean 45-year changes estimated for each of the 37 bias-corrected and downscaled CMIP6 simulations for the preindustrial (1860–1904) and modern (1980–2024) periods, as described above.

For each climate period, we counted the number of simulations with a trend at least as large as the observed GridMET trend. For season length and last day, this corresponded to trends equal to or greater than the observed positive trend. For first day, for which negative values indicate earlier onset, we counted trends equal to or more negative than the observed trend. We calculated the empirical probability of observing a trend at least as large as the GridMET trend as (Eq. 2):

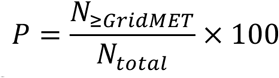

where *N*_≥_*_GridMET_* is the number of simulations meeting or exceeding the observed trend and *N_total_* = 37. We additionally calculated the proportion of simulations with a trend in the same direction as the observed trend, defined as trends ≥0 for season length and last day and trends ≤0 for first day.

We report one-sided 95% upper confidence bounds for all empirical probabilities because season length and last day in the preindustrial scenario had zero exceedances. A point estimate of zero would incorrectly suggest that there is a 0% probability of observing a trend at least as large as the observed trend under preindustrial conditions. With only 37 climate simulations, zero exceedances cannot distinguish a true probability of zero from a small nonzero probability. We therefore used one-sided 95% upper confidence bounds to account for uncertainty associated with the finite ensemble size. Bounds were calculated as exact (Clopper–Pearson) binomial bounds. For *^K^* exceedances among *^N^* simulations, the upper bound is the value *^X^* satisfying (Eq. 3):

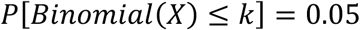

When *k* = 0, this reduces to:

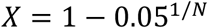

or 7.8% for *N* = 37. Thus, zero exceedances do not imply that the true probability is zero; rather, they indicate that the probability is sufficiently small that zero exceedances would be expected in a finite ensemble. For *N* = 37, a true probability greater than 7.8% would result in at least one exceedance with greater than 95% probability. We therefore interpret 7.8% as the one-sided 95% upper bound when zero exceedances are observed. We calculated the 95% upper bound for all scenarios, including those with one or more exceedances, to standardize comparisons across climate periods and season characteristics.

To quantify the change in the probability of observing trends at least as large as those observed under modern relative to preindustrial climate conditions, we calculated the ratio of the corresponding one-sided 95% upper confidence bounds (Eq.4):

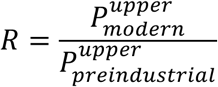

where 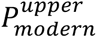 and 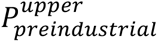 are the one-sided 95% upper confidence bounds for the modern (1980–2024) and preindustrial (1860–1904) periods, respectively. We used the upper confidence bounds rather than point estimates so that the ratio remained defined when no preindustrial simulations met or exceeded the observed trend. This ratio describes the relative rarity of the observed trend under modern versus preindustrial climate conditions and does not represent the probability that anthropogenic climate change caused the observed trend.

## Results

### Changes in temperature-based transmission season length

To assess temperature effects on WNV transmission season length, temperature data were filtered by days ≥16.7°C (See Methods), which falls within the thermal suitability for WNV transmission by *Cx. pipiens* populations in NYS^16^. Hereafter, we refer to the number of days above the thermal suitability threshold as the ‘transmission season’. For all transmission season metrics, the best-fitting models included quadratic effects of year that interacted with elevation, demonstrating that temporal changes in season timing and duration differed across the elevation gradient, with the largest increases occurring in lower-elevation counties (Table 1; Tables S1–S3). The average transmission season length has significantly increased over the last few decades (Fig. 1A; Table 1). Although the increase varies by county (Fig. 1DG), the transmission season at the average elevation has extended by 29.6 days (95% credible interval (CrI): 26.4, 32.5) from 1980 to 2024, and by 19.7 days (95% CrI: 16.4, 23.1) since 1999-2024 (Table 1). This expansion was greatest in low-elevation and high mean temperature counties: for instance, Kings County (average elevation: 9 m and average temperature: 13.29°C) gained 27.1 days from 1999 to 2024 (34.9 days since 1980), compared to 10.8 days (23.2 days since 1980) in Hamilton County, the highest-elevation county in the state (average elevation: 612 m and average temperature: 5.38°C) (Fig. 1DG; Table 1).

**Figure 1.**
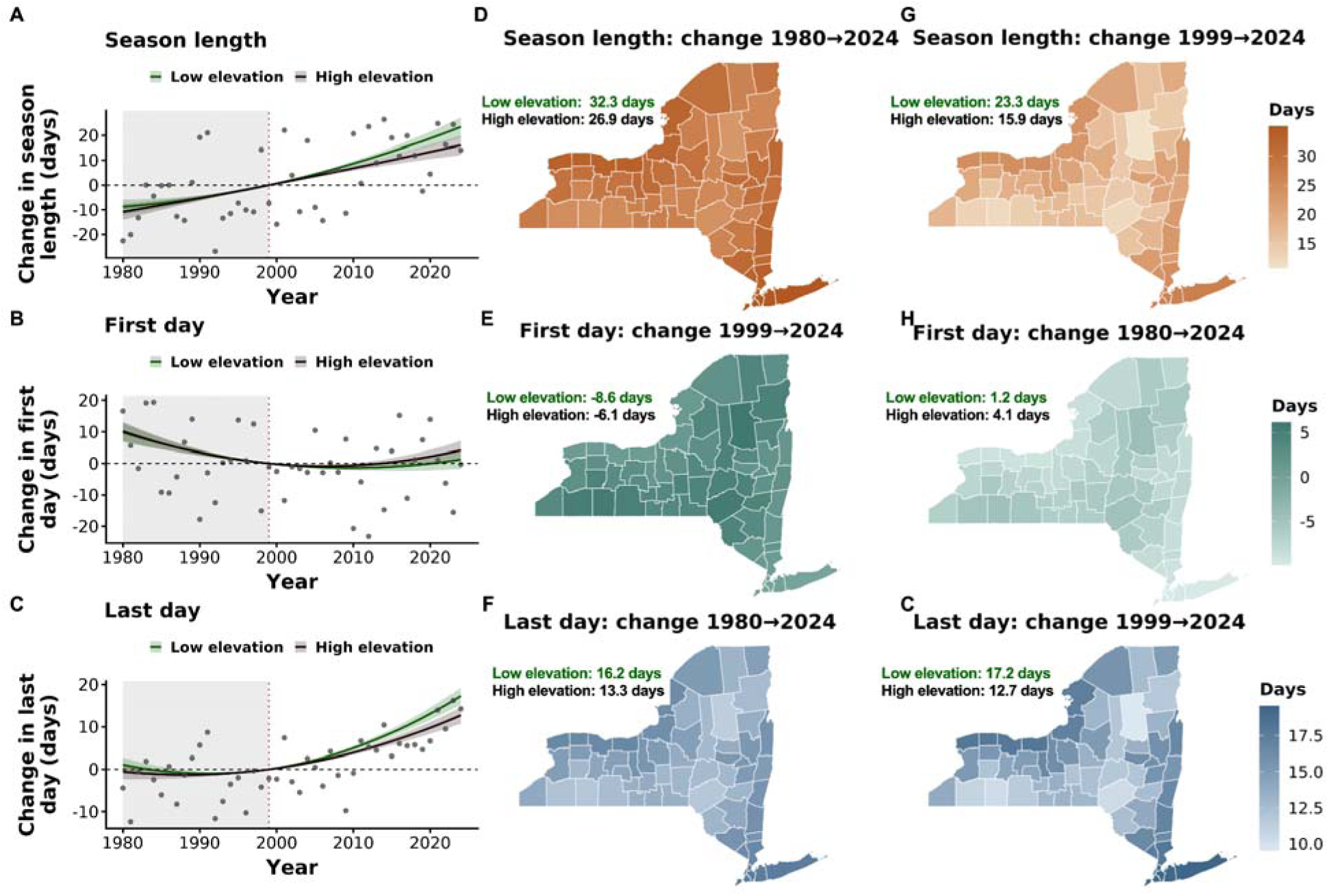
Estimated changes in West Nile virus transmission season length, start date, and end date across New York State counties from 1980–2024, modeled using a Bayesian spatiotemporal framework. (A-C) Posterior mean trends relative to 1999 (WNV emergence year) for low-(mean: 146 m; blue) and high-elevation (mean: 458 m; red) county groups (median split at 265 m); shaded ribbons = 95% credible intervals. **(D-I)** county-level posterior mean change from 1980–2024 and 1999–2024, respectively; map subtitles report group-level posterior means and 95% credible intervals.

**Table 1.** Model coefficients and mean estimates for each Bayesian spatiotemporal hierarchical model. Coefficients represent days per unit of the explanatory variables (e.g., year = days per year). Means are based on state-wide average elevation. Values in parentheses indicate the 95% credible intervals. Coefficient estimates listed in bold indicate variables of interest for which credible intervals did not overlap zero.

| Term | Season Length | First Day | Last Day |
| --- | --- | --- | --- |
| Intercept | -0.78 (-1.91, 0.35) | -2.26 (-3.26, -1.26) | -2.29 (-2.88, -1.69) |
| Year (linear) | **0.67 (0.60, 0.74)** | -0.17 (-0.22, -0.12) | **0.33 (0.30, 0.37)** |
| Year<sup>2</sup> (quadratic) | **0.006 (0.001, 0.011)** | **0.014 (0.009, 0.019)** | **0.014 (0.011, 0.017)** |
| Elevation | -7.15 (-9.07, -5.25) | **4.03 (2.68, 5.39)** | -3.08 (-3.98, -2.18) |
| Year × Elevation | -0.073 (-0.123, -0.022) | 0.034 (-0.007, 0.075) | -0.038 (-0.065, -0.012) |
| Year<sup>2</sup> × Elevation | -0.005 (-0.010, -0.001) | 0.002 (-0.002, 0.005) | -0.004 (-0.006, -0.001) |
| Est. change, 1999–2024 (days) | 19.7 (16.4, 23.1) | 2.7 (0.1, 5.2) | 15.0 (13.3, 16.7) |
| Est. change, 1980–2024 (days) | 29.6 (26.4, 32.5) | -7.3 (-9.5, -5.2) | 14.7 (13.0, 16.3) |

When considering conditions leading up to WNV emergence, the season start date shifted earlier by approximately 7.3 days (95% CrI: -9.5, -5.2) at average elevation over the full 1980–2024 period (Fig. 1BEH; Table 1). However, most of this advance preceded WNV emergence in New York: after 1999, changes in season start date were small and statistically uncertain, with the credible interval close to 0 (2.6 days; 95% CrI: 0.1, 5.2; Fig. 1BEH). In contrast, season end date extended substantially and continued to do so after 1999, with nearly the entire 1980–2024 shift occurring in the post-emergence period (15.0 days, 95% CrI: 13.3, 16.7) (Fig. 1CFI; Table 1). Model estimates for season length and last day showed little sensitivity to the threshold used (Fig. S2). First day was more sensitive: higher temperature thresholds derived from the upper credible interval of temperature-minima estimates yielded significantly earlier season start dates between 1999–2024, particularly when using population-specific temperature-minima (Fig. S2).

### Relationship between season length and epidemiological outcomes

Change in season length (incorporated as a smoothed function) was a significant nonlinear predictor of total WNV incidence from 1999-2024 (Fig. 2A; Table 2). By contrast, water availability (average summer SPEI) did not have a significant effect (Fig. 2A; Table 2). Counties with the greatest season lengthening tended to have the highest incidence, with the effect accelerating at higher values. Overall, the generalized additive model, which included only fixed effects, explained 46.4% of the deviance.

**Figure 2.**
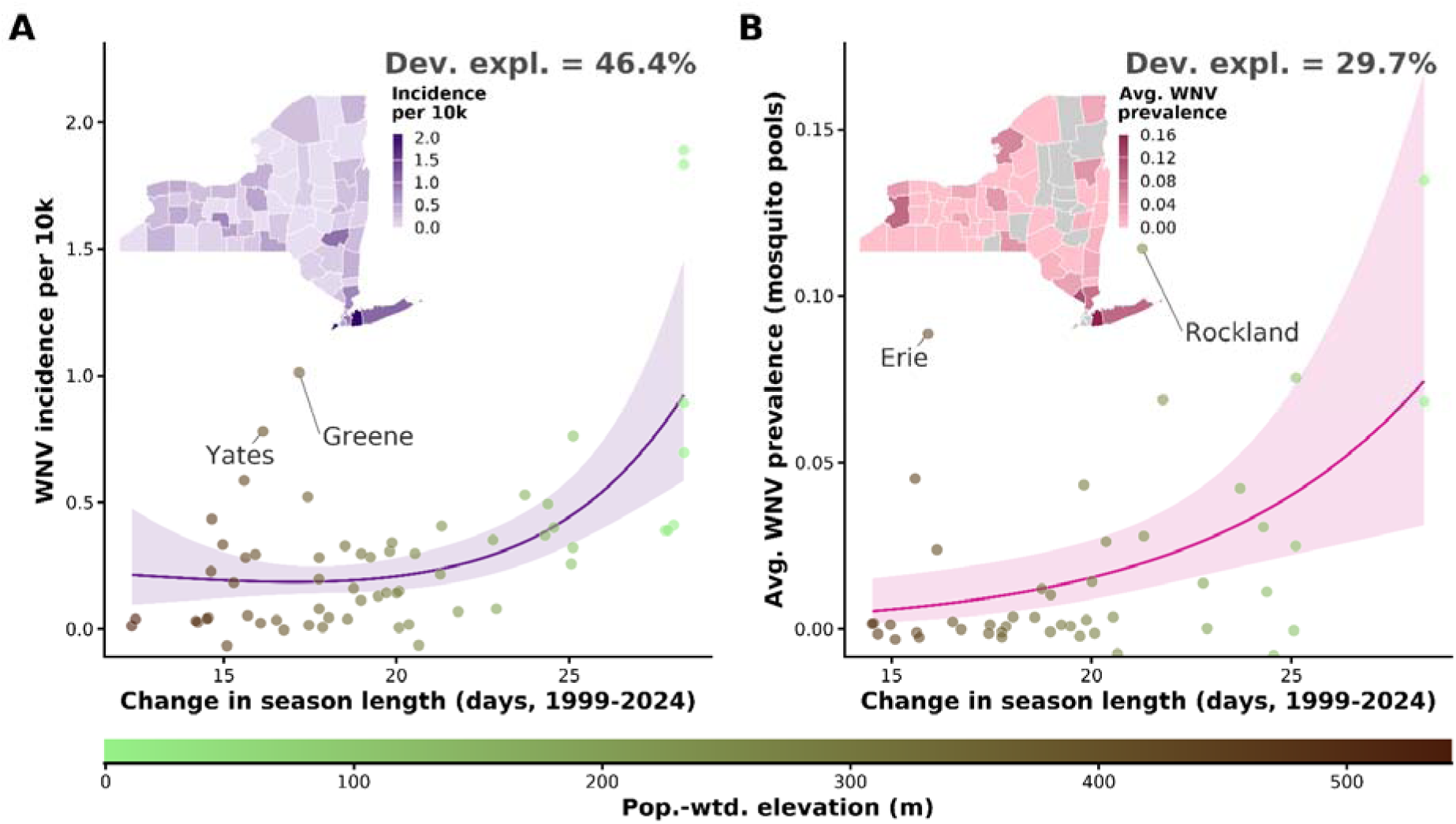
Association between change in WNV transmission season length and WNV across New York State counties from 1999–2024. **A**) Cumulative WNV human case incidence (per 10,000 population) versus estimated change in transmission season length (days, 1999–2024). **B**) Average annual WNV mosquito pool prevalence versus estimated change in transmission season length. In both panels, points show partial residuals (i.e., the observed outcome adjusted for average summer SPEI-90d) and are colored by county population-weighted elevation. Smooth lines show generalized additive model (GAM) fits with 95% confidence ribbons, evaluated at mean summer SPEI-90d. Deviance explained by the season length smooth is annotated in each panel. Inset maps show the spatial distribution of cumulative WNV human case incidence (**A**) and average mosquito pool prevalence (**B**) across New York State counties; grey areas on the map indicate counties without mosquito prevalence data. Season length changes are county-specific estimates derived from the best-fitting INLA model at each county’s population-weighted elevation.

**Table 2.** Generalized additive model results for WNV human case incidence and mosquito pool prevalence in New York State counties, 1999–2024. Estimates for parametric terms are on the link scale (log for incidence, logit for prevalence). Season length change is the estimated change in WNV transmission season length (days) from 1999 to 2024, derived from the final season length INLA models at each county’s population-weighted elevation. Avg. summer SPEI = mean SPEI-90d across June–August, 1999–2024. The incidence model includes log(mean population) as an offset. Smooth term edf = estimated degrees of freedom; test statistics differ by family (χ² for negative binomial, F for quasibinomial). Coefficients listed in bold have credible intervals that do not overlap zero.

| Term | Estimate (95% CI) | Test statistic | *p* |
| --- | --- | --- | --- |
| **WNV Incidence** |  |  |  |
| *Linear terms* |  |  |  |
| Intercept | −10.45 (−10.72, −10.17) | **$z = -74.7$** | **<0.001** |
| Avg. summer SPEI | −1.73 (−4.42, 0.97) | $z = -1.26$ | 0.208 |
| *Smooth terms* |  |  |  |
| s(Season length change) | **edf = 1.83** | **$\chi^2 = 32.0$** | **<0.001** |
| *Model fit* |  |  |  |
| Deviance explained | 46.40% |  |  |
| *n* (counties) | 62 |  |  |
| Family | Negative binomial |  |  |
| **WNV Prevalence** |  |  |  |
| *Linear terms* |  |  |  |
| Intercept | −4.06 (−4.70, −3.42) | **$t = -12.5$** | **<0.001** |
| Avg. summer SPEI | −2.58 (−8.54, 3.37) | $t = -0.85$ | 0.4 |
| *Smooth terms* |  |  |  |
| s(Season length change) | **edf = 1.00** | **$F = 9.72$** | **0.003** |
| *Model fit* |  |  |  |
| Deviance explained | 29.70% |  |  |
| *n* (counties) | 46 |  |  |
| Family | Quasibinomial |  |  |

Season length change was also a significant predictor of average WNV prevalence in mosquito pools (Fig. 2B; Table 2). The relationship was approximately linear (effective degrees of freedom ≈ 1.0; effective degrees of freedom measures complexity of the smooth term), indicating a constant positive relationship between season expansion and vector infection rates across the 46 counties with prevalence data. Average summer SPEI was again non-significant (Fig. 2B; Table 2). Overall, the model explained a moderate proportion of deviance (29.7%).

Moran’s I tests on GAM residuals detected no significant spatial autocorrelation in either the incidence model (I = −0.061; p = 0.734) or the prevalence model (I = 0.023; p = 0.285), supporting the assumption of spatially independent errors after conditioning on season length change and SPEI and justifying the use of a generalized additive model as opposed to generalized additive mixed model with random effects. Further, the effect of season length remained significant and positive for all leave-one-out model tests, indicating that no one county was driving the significant effect. Deviance explained ranged from 39.2% to 51.8% across iterations for the incidence models and 16.2% to 40.1% for the prevalence models (Fig. S3). Nassau County, which had both the largest season length change and highest mosquito pool prevalence, was the most influential observation in the prevalence model; its exclusion reduced deviance explained from 29.7% to 16.2%, though the association between season length change and prevalence remained statistically significant (p = 0.026).

### Climate change attribution analysis

To investigate the sensitivity of the duration of the WNV transmission season to historical climate forcing, we analyzed 37 climate model simulations from the CMIP6 archive. When analyzing WNV season characteristics in each of these climate model simulations during the recent period relative to the historical baseline used to approximate climate conditions prior to major anthropogenic forcing (1860-1904)^45^, we found robust changes in WNV season length, onset date, and termination date (Figs. 3 and S4-6). We also found statistically significant (Wilcoxon signed rank test; p < 0.001 for all metrics) changes in the distribution of WNV season length, first day, and last day over the last 45 years (1980-2024) of the climate model simulations compared to the preindustrial period (1860-1904) (Figs. 3 and S4-6; Table S4). In particular, we found that on average across climate model simulations and all NYS counties, WNV season length increased by ∼22.4 days (100% of values > 0; Fig. 3A), onset date decreased by ∼10.9 days (97.3% of values < 0; Fig 3B), and termination date increased by ∼9.7 days (94.6% of values > 0;Fig. 3C).

**Figure 3.**
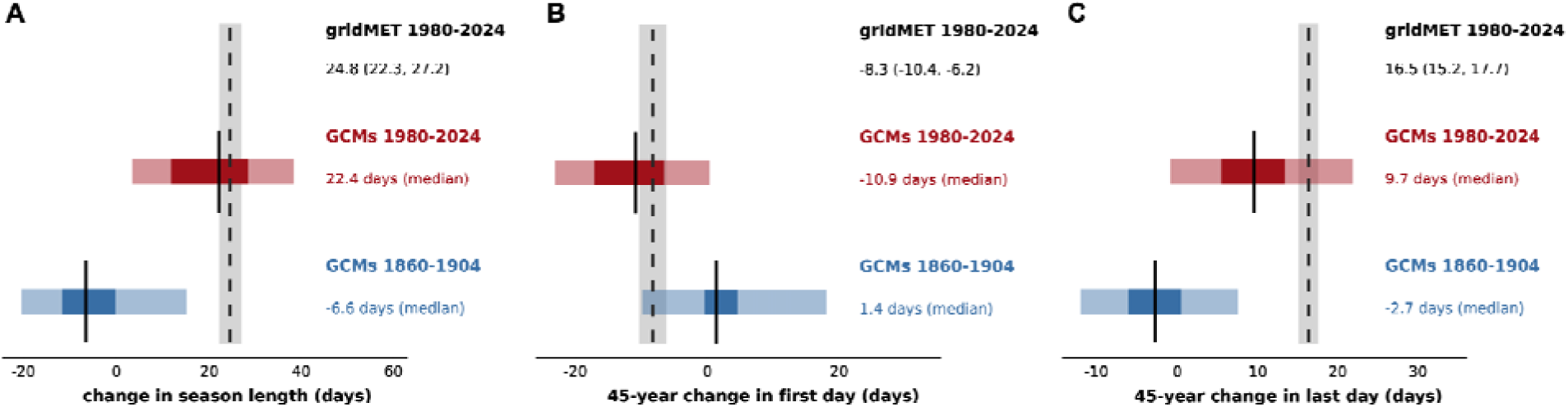
Influence of global warming on trends in the seasonality of WNV transmission. Modeled changes in WNV transmission season length under two 45-year windows: 1980–2024 (modern/observational era) and 1860–1904 (pre-industrial-era analog), using an ensemble of 37 GCMs. **(A–C)** Distributions of the 45-year change in (**A**) season length (days), (**B**) first day, and (**C**) last day of the transmission season estimated by the GCM ensemble for each period. Dark and light shading indicate the interquartile and 2.5%-97.5% quantile ranges, respectively, across the 37 models; the solid black line indicates the ensemble median. The black dashed line and shaded band indicate the observed change in season length (posterior mean and 95% credible interval) estimated from GridMET climate data over 1980–2024. The GridMET estimate is shown for comparison with the GCM ensemble and represents the posterior mean and 95% credible interval from the model, whereas GCM estimates represent the median change across climate simulations.

Although we cannot independently observe the influence of global warming on the recent observed trends in WNV season characteristics, we can analyze trends in the climate model simulations to determine whether global warming increased the likelihood of the observed trends in season length, onset, and termination. We compared the GridMET trends to the distribution of 45-year trends in the climate model simulations calculated from the preindustrial period (1860-1904) and the recent period (1980-2024) (Figs. 3 and S4-6). We found that the probability of trends in WNV season length as extreme as the GridMET trend, which was used to measure the observed trend, increased from 0% (CI: 0%, 9.5%) in the pre-industrial period to 40.5% (CI: 24.8%, 57.9%) in the recent period (Fig. S4; Table S5).

In order to quantitatively compare the likelihood of the observed trend in the preindustrial and recent periods, we used an exact binomial approach to estimate the probability of model-simulated trends at least as large as the observed GridMET trend and compared these probabilities between climate periods. We conservatively estimate the observed trend was at least 7.1 times as likely to occur due to global warming, using the upper bound of the pre-industrial probability’s 95% confidence interval as a conservative denominator to account for the effects of sampling error caused by our relatively small sample size (N=37)(Figs. 3 and S4; Table S5). Similarly, we found that global warming since 1850 has increased the likelihood of the observed trends in season onset and termination dates, implying that the observed trend in WNV season start date was 4.7 times as likely to occur because of global warming, and that the observed trend in WNV season end date was at least 3.4 times as likely (Figs. 3 and S5-6; Table S5).

## Discussion

Temperature is a key driver of vector-borne pathogen transmission, yet its influence on transmission season length remains poorly characterized for many diseases, including WNV. Using county-level temperature data and the lower thermal limit for WNV transmission in *Cx. pipiens*, our best-fitting model indicated that transmission season length increased at an accelerating rate over time, with the highest magnitude and rate of expansion at lower elevations. We show that the WNV transmission season in NYS lengthened by an average of 29.6 days over 1980–2024, and by 19.7 days over 1999–2024, the period during which WNV emerged and spread throughout the state (Fig. 1A), with expansion varying by county and greatest in low-elevation counties (Fig. 1D–G). Thermally suitable transmission seasons began earlier and terminated later, with larger shifts at the end of the season (Fig. 1B–C). Bayesian spatiotemporal hierarchical modeling further showed that longer transmission seasons are associated with increased WNV incidence in humans (Fig. 2A) and higher WNV prevalence in mosquito surveillance pools (Fig. 2B). Climate model simulations indicate that anthropogenic greenhouse gas emissions have substantially contributed to these changes. It is possible but highly unlikely (<5% probability) that the observed increase in WNV transmission season length would have occurred in the absence of global warming since 1900 (Fig. 3). Further, observed trends in transmission season length, season onset, and season termination were approximately 7.1, 4.7, and 3.4 times as likely, respectively, under present-day climate conditions than under counterfactual scenarios without anthropogenic warming (Figs. 3 and S4-6). Together, these results demonstrate that climate change has lengthened the WNV transmission season in NYS, with measurable consequences for vector infection dynamics and human disease risk.

Although the observed expansion in transmission season is generally consistent with warming-driven increases in thermal suitability, this pattern is unlikely to remain spatially uniform in the future. Continued climate warming may differentially affect counties depending on baseline temperature conditions, potentially altering the magnitude and direction of seasonal shifts. Vector-borne diseases are highly sensitive to warming-driven extensions of the transmission season, which may modify thermal suitability windows, increase opportunities for vector expansion, and facilitate evolutionary responses in both vectors and viruses^13,46–48^. However, sustained warming could eventually reduce transmission suitability in some regions in the hottest portion of the season if temperatures exceed upper thermal limits, although this is not expected in New York in the near term^14^.

Beyond spatial heterogeneity in climate exposure, GAMs revealed variability in how season length change relates to WNV outcomes across counties. Associations between WNV incidence and season length change were heterogeneous, with Greene and Yates Counties –– both historically rural and with an average elevation of 1100-1200 ft –– exhibiting elevated incidence relative to modeled expectations. However, both counties had very small populations and few reported cases, which likely inflated incidence estimates. Similarly, mosquito pool prevalence varied substantially across counties, with urban counties like Nassau, Rockland, and Erie showing the highest observed prevalence relative to model predictions, potentially reflecting more consistent and long-term surveillance efforts in these regions, as well as environmental conditions favorable for mosquito populations, including proximity to major water bodies and associated wetland habitats.

WNV incidence estimates were less sensitive to individual county-level contributions than mosquito pool prevalence (Fig. S3), suggesting greater robustness in human incidence patterns compared to surveillance-derived prevalence metrics. This difference likely reflects variation in mosquito sampling intensity and reporting structure across counties, which can introduce additional uncertainty into prevalence-based estimates. However, the relationships between season length expansion and both human cases and mosquito prevalence were robust to exclusion of individual counties, suggesting general relationships.

While temperature is a primary driver of WNV seasonality, other climatic factors—including precipitation and humidity—also shape transmission dynamics^6,8,21,49^. Consistent with previous studies in Europe and the US, warmer conditions are associated with extended transmission seasons and increased WNV activity seasonality^10,21,50–52^. However, disentangling the relative contribution of correlated environmental drivers remains an important challenge for future work, particularly under non-stationary climate conditions.

Our analysis is subject to several limitations. First, mosquito surveillance data do not include New York City (NYC), where WNV was initially detected in the state and where a substantial proportion of human cases occur. While this omission does not bias county-level associations, inclusion of NYC data could refine estimates of prevalence and transmission intensity. Second, improved resolution of human case detection and reporting could further clarify the relationship between seasonal length and disease burden. Finally, although avian hosts are central to WNV transmission dynamics^53–56^, climate-driven changes in bird community composition could not be incorporated due to limited surveillance data.

Our results have important implications for optimizing WNV surveillance and control by supporting the initiation of control measures earlier in the spring and extending them later into the fall. Expanding the seasonal window for surveillance and intervention may help prevent early transmission and the establishment of infections while improving estimates of WNV prevalence and incidence later in the season. In addition to enhancing vector control efforts, these findings may justify increased investment in surveillance and diagnostic testing of mosquito populations and dead-end hosts, including humans and horses, thereby improving early detection and public health response.

Together, these findings highlight WNV as a clear example of how climate-driven seasonal expansions can amplify vector-borne diseases. Our results indicate that recent warming has extended the temporal window for transmission in NYS, with implications for both mosquito infection dynamics and human disease risk. As climate change continues in the coming decades^57^, similar shifts in thermal suitability, vector activity, and host–vector interactions are likely to influence the distribution, intensity, and timing of WNV and other vector-borne diseases.

## Acknowledgments

We are grateful to the New York State Arbovirology Laboratory, particularly Elyse Banker and Alexis Russell, for their support throughout this study. We thank the New York State Department of Health for providing human WNV case data and the New York State Bureau of Communicable Disease Control and county health departments for coordinating and implementing mosquito surveillance collections, especially Jennifer White. We also thank Dr. Alexander Keyel for his valuable assistance and insightful feedback. We acknowledge the World Climate Research Programme, which coordinated and promoted CMIP6, and thank the climate modeling groups for producing and making available their model output, the Earth System Grid Federation (ESGF) for archiving the data and providing access, and the multiple funding agencies that support CMIP6 and ESGF.

## Contributions

RLF and EAM conceived and designed the study. RLF, CKG, JTT, and ATC collected and formatted the data utilized in the manuscript. RLF, CKG, and JTT conducted the results analysis. RLF, CKG, and JTT prepared the manuscript. NSD and EAM supervised the work. CKG, JTT, NSD, ATC, and EAM provided constructive comments to improve the manuscript and participated in revising the manuscript. Code utilized for temperature and host data analysis can be found at https://github.com/Rachellfay/WNV-Season-Length.

## Declaration of interests

None.

## Data Sharing

Due to confidentiality restrictions set by the New York State Department of Health (NYSDOH), mosquito surveillance data cannot be made publicly available. However, these data can be requested directly from the NYSDOH Bureau of Communicable Diseases at. Human case data were downloaded from the National Center for Emerging and Zoonotic Infectious Diseases https://www.cdc.gov/west-nile-virus/data-maps/historic-data.html. Model output from Phase 6 of the Coupled Model Intercomparison Project is available from the Earth System Grid Federation through their website at https://esgf.github.io/. The GridMET data used in this analysis is available from their website at https://www.climatologylab.org/. The New York State County boundaries used in this analysis can be downloaded from https://www.ny.gov/.

## Notes

https://github.com/Rachellfay/WNV-Season-Length

### Competing Interest Statement

The authors have declared no competing interest.

### Summary of Updates

We have updated our climate attribution model, which yielded new results.

