## Supplemental Figures for "The impact of climate change on transmission season length: West Nile virus as a case study"

**Supplementary figures and tables**

**Table S1. Comparison of Bayesian spatiotemporal INLA models predicting annual WNV transmission season length in NYS.** Models differed in their temporal trend specification (linear or quadratic year), spatial covariates (elevation, latitude, or county-specific effects), and climate covariates (no ENSO term, linear ENSO effect, binary El Niño indicator, or a second-order random walk [RW2] representation of ENSO). Model performance was evaluated using the Deviance Information Criterion (DIC), effective number of parameters (pD), Widely Applicable Information Criterion (WAIC), and the log Conditional Predictive Ordinate (LCPO). Lower DIC and WAIC values indicate better model fit, while higher (less negative) LCPO values indicate better predictive performance. The estimated statewide change in transmission season length from 1999 to 2024 is reported as the posterior median (total_est) with 95% credible intervals (total_lo–total_hi). ΔDIC, ΔWAIC, and ΔLCPO represent differences relative to the best-performing model (M12).

| **model** | **DIC** | **pD** | **WAIC** | **LCPO** | **total_est** | **total_lo** | **total_hi** | **dDIC** | **dWAIC** | **dLCPO** |
| --- | --- | --- | --- | --- | --- | --- | --- | --- | --- | --- |
| M12: quadratic year x elev + ENSO RW2 | 5920.827 | 65.698 | 5920.965 | -2960.521 | 19.667 | 15.399 | 23.938 | 0.000 | 0.000 | 0.000 |
| M19: linear year x elevation + ENSO RW2 | 5926.744 | 63.879 | 5927.058 | -2963.564 | 16.376 | 14.596 | 18.158 | 5.917 | 6.093 | -3.044 |
| M16: quadratic year x lat + ENSO RW2 | 5945.299 | 75.890 | 5945.456 | -2972.783 | 19.667 | 15.387 | 23.950 | 24.472 | 24.491 | -12.263 |
| M4: quadratic year + ENSO RW2 | 5949.925 | 73.672 | 5950.511 | -2975.307 | 19.667 | 15.384 | 23.953 | 29.098 | 29.547 | -14.786 |
| M17: linear year + ENSO RW2 | 5951.754 | 73.107 | 5952.217 | -2976.158 | 16.371 | 14.589 | 18.156 | 30.927 | 31.252 | -15.637 |
| M20: linear year x latitude + ENSO RW2 | 5952.468 | 73.368 | 5953.040 | -2976.572 | 16.374 | 14.587 | 18.163 | 31.641 | 32.076 | -16.051 |
| M8: quadratic year + county + ENSO RW2 | 5957.801 | 86.113 | 5957.045 | -2978.608 | 19.667 | 15.276 | 24.061 | 36.974 | 36.080 | -18.088 |
| M18: linear year x county + ENSO RW2 | 5959.626 | 85.553 | 5958.564 | -2979.364 | 16.374 | 14.479 | 18.271 | 38.799 | 37.599 | -18.844 |
| M26: quadratic year x elev + El Nino binary | 6645.354 | 45.846 | 6644.841 | -3322.435 | 19.510 | 15.738 | 23.282 | 724.527 | 723.876 | -361.914 |
| M25: linear year x elev + El Nino binary | 6665.045 | 43.589 | 6664.988 | -3332.507 | 14.314 | 12.876 | 15.752 | 744.218 | 744.023 | -371.987 |
| M28: quadratic year x lat + El Nino binary | 6670.861 | 56.766 | 6670.469 | -3335.258 | 19.510 | 15.726 | 23.293 | 750.034 | 749.504 | -374.737 |
| M22: quadratic year + El Nino binary | 6673.156 | 54.492 | 6673.355 | -3336.699 | 19.510 | 15.724 | 23.295 | 752.329 | 752.391 | -376.178 |
| M24: quadratic year + county + El Nino binary | 6681.474 | 63.474 | 6680.616 | -3340.345 | 19.510 | 15.603 | 23.416 | 760.647 | 759.651 | -379.825 |
| M21: linear year + El Nino binary | 6688.966 | 52.202 | 6689.310 | -3344.675 | 14.314 | 12.874 | 15.755 | 768.139 | 768.346 | -384.154 |
| M27: linear year x lat + El Nino binary | 6691.845 | 54.633 | 6691.952 | -3345.997 | 14.314 | 12.873 | 15.755 | 771.018 | 770.987 | -385.477 |
| M10: quadratic year x elevation | 6698.957 | 44.510 | 6698.479 | -3349.253 | 19.543 | 15.736 | 23.350 | 778.130 | 777.515 | -388.733 |
| M23: linear year x county + El Nino binary | 6699.293 | 62.950 | 6698.468 | -3349.270 | 14.314 | 12.741 | 15.887 | 778.466 | 777.504 | -388.750 |
| M11: linear year x elevation + ENSO linear | 6709.545 | 43.453 | 6709.255 | -3354.640 | 14.221 | 12.772 | 15.670 | 788.718 | 788.291 | -394.120 |
| M9: linear year x elevation | 6720.332 | 42.422 | 6720.195 | -3360.110 | 14.066 | 12.616 | 15.516 | 799.505 | 799.230 | -399.589 |
| M14: quadratic year x latitude | 6724.637 | 55.536 | 6724.268 | -3362.157 | 19.543 | 15.725 | 23.361 | 803.810 | 803.304 | -401.636 |
| M2: quadratic year | 6726.753 | 53.228 | 6726.932 | -3363.486 | 19.543 | 15.724 | 23.362 | 805.926 | 805.968 | -402.966 |
| M3: linear year + ENSO linear | 6734.830 | 53.197 | 6734.996 | -3367.518 | 14.221 | 12.767 | 15.675 | 814.003 | 814.031 | -406.998 |
| M6: quadratic year + county slope | 6736.028 | 62.920 | 6735.415 | -3367.745 | 19.543 | 15.589 | 23.497 | 815.201 | 814.451 | -407.224 |
| M15: linear year x latitude + ENSO linear | 6736.477 | 54.435 | 6736.372 | -3368.207 | 14.221 | 12.767 | 15.675 | 815.650 | 815.407 | -407.687 |
| M7: linear year x county + ENSO linear | 6744.328 | 63.256 | 6743.301 | -3371.687 | 14.221 | 12.634 | 15.808 | 823.501 | 822.336 | -411.166 |
| M1: linear year | 6745.587 | 52.254 | 6745.885 | -3372.962 | 14.066 | 12.612 | 15.520 | 824.760 | 824.920 | -412.441 |
| M13: linear year x latitude | 6747.219 | 53.429 | 6747.345 | -3373.693 | 14.066 | 12.612 | 15.520 | 826.392 | 826.381 | -413.172 |
| M5: linear year x county slope | 6754.595 | 61.320 | 6754.069 | -3377.070 | 14.066 | 12.479 | 15.653 | 833.768 | 833.104 | -416.549 |

**Table S2. Comparison of Bayesian spatiotemporal INLA models predicting the first day of the WNV transmission season in NYS.** Models differed in their temporal trend specification (linear or quadratic year), spatial covariates (elevation, latitude, or county-specific effects), and climate covariates (no ENSO term, linear ENSO effect, binary El Niño indicator, or a second-order random walk [RW2] representation of ENSO). Model performance was evaluated using the Deviance Information Criterion (DIC), effective number of parameters (pD), Widely Applicable Information Criterion (WAIC), and the log Conditional Predictive Ordinate (LCPO). Lower DIC and WAIC values indicate better model fit, while higher (less negative) LCPO values indicate better predictive performance. The estimated statewide change in the first day of the transmission season from 1999 to 2024 is reported as the posterior median (total_est) with 95% credible intervals (total_lo–total_hi). Negative estimates indicate an earlier onset of the transmission season, whereas positive estimates indicate a later onset. ΔDIC, ΔWAIC, and ΔLCPO represent differences relative to the best-performing model (M12).

| **model** | **DIC** | **pD** | **WAIC** | **LCPO** | **total_est** | **total_lo** | **total_hi** | **dDIC** | **dWAIC** | **dLCPO** |
| --- | --- | --- | --- | --- | --- | --- | --- | --- | --- | --- |
| M12: quadratic year x elev + ENSO RW2 | 6677.586 | 60.206 | 6676.899 | -3338.479 | 2.623 | -0.977 | 6.223 | 0.000 | 0.000 | 0.000 |
| M16: quadratic year x lat + ENSO RW2 | 6695.457 | 69.771 | 6694.759 | -3347.423 | 2.623 | -0.982 | 6.228 | 17.872 | 17.861 | -8.944 |
| M4: quadratic year + ENSO RW2 | 6697.602 | 67.259 | 6697.628 | -3348.854 | 2.615 | -0.989 | 6.219 | 20.016 | 20.729 | -10.374 |
| M8: quadratic year + county + ENSO RW2 | 6708.598 | 75.191 | 6708.743 | -3354.432 | 2.620 | -1.058 | 6.299 | 31.012 | 31.844 | -15.952 |
| M19: linear year x elevation + ENSO RW2 | 6724.919 | 67.116 | 6724.931 | -3362.504 | -3.926 | -5.157 | -2.696 | 47.333 | 48.033 | -24.024 |
| M17: linear year + ENSO RW2 | 6730.446 | 65.904 | 6730.357 | -3365.216 | -3.932 | -5.166 | -2.698 | 52.860 | 53.458 | -26.737 |
| M20: linear year x latitude + ENSO RW2 | 6731.431 | 65.597 | 6731.163 | -3365.619 | -3.929 | -5.163 | -2.695 | 53.845 | 54.264 | -27.139 |
| M18: linear year x county + ENSO RW2 | 6740.576 | 73.340 | 6740.687 | -3370.399 | -3.932 | -5.235 | -2.629 | 62.990 | 63.788 | -31.920 |
| M11: linear year x elevation + ENSO linear | 7487.030 | 35.940 | 7486.315 | -3743.166 | -4.871 | -6.061 | -3.681 | 809.444 | 809.416 | -404.686 |
| M26: quadratic year x elev + El Nino binary | 7497.400 | 38.658 | 7496.389 | -3748.205 | -1.477 | -4.648 | 1.693 | 819.814 | 819.490 | -409.726 |
| M10: quadratic year x elevation | 7505.358 | 37.493 | 7504.316 | -3752.168 | -1.574 | -4.718 | 1.571 | 827.772 | 827.418 | -413.689 |
| M25: linear year x elev + El Nino binary | 7507.005 | 36.973 | 7506.459 | -3753.239 | -4.956 | -6.163 | -3.750 | 829.420 | 829.560 | -414.760 |
| M3: linear year + ENSO linear | 7510.037 | 44.521 | 7509.652 | -3754.840 | -4.871 | -6.065 | -3.677 | 832.451 | 832.753 | -416.361 |
| M15: linear year x latitude + ENSO linear | 7511.871 | 44.916 | 7510.831 | -3755.429 | -4.871 | -6.065 | -3.677 | 834.286 | 833.932 | -416.949 |
| M9: linear year x elevation | 7512.116 | 35.534 | 7511.489 | -3755.753 | -4.700 | -5.895 | -3.506 | 834.531 | 834.590 | -417.274 |
| M28: quadratic year x lat + El Nino binary | 7517.190 | 47.449 | 7516.235 | -3758.134 | -1.477 | -4.653 | 1.698 | 839.604 | 839.336 | -419.654 |
| M22: quadratic year + El Nino binary | 7517.519 | 45.476 | 7517.222 | -3758.626 | -1.477 | -4.653 | 1.699 | 839.933 | 840.324 | -420.147 |
| M7: linear year x county + ENSO linear | 7520.960 | 51.621 | 7519.278 | -3759.660 | -4.871 | -6.147 | -3.595 | 843.374 | 842.379 | -421.181 |
| M14: quadratic year x latitude | 7525.129 | 46.053 | 7524.271 | -3762.151 | -1.574 | -4.726 | 1.579 | 847.543 | 847.372 | -423.672 |
| M2: quadratic year | 7525.697 | 44.471 | 7525.354 | -3762.691 | -1.574 | -4.725 | 1.578 | 848.111 | 848.455 | -424.212 |
| M24: quadratic year + county + El Nino binary | 7527.622 | 52.031 | 7525.979 | -3763.012 | -1.477 | -4.729 | 1.775 | 850.037 | 849.080 | -424.532 |
| M21: linear year + El Nino binary | 7527.999 | 44.454 | 7527.902 | -3763.965 | -4.956 | -6.166 | -3.747 | 850.413 | 851.003 | -425.486 |
| M27: linear year x lat + El Nino binary | 7531.077 | 45.500 | 7530.727 | -3765.379 | -4.956 | -6.167 | -3.746 | 853.492 | 853.828 | -426.899 |
| M1: linear year | 7533.586 | 43.368 | 7533.417 | -3766.722 | -4.700 | -5.897 | -3.503 | 856.001 | 856.518 | -428.242 |
| M6: quadratic year + county slope | 7535.798 | 51.039 | 7534.386 | -3767.214 | -1.574 | -4.794 | 1.647 | 858.212 | 857.487 | -428.734 |
| M13: linear year x latitude | 7536.615 | 44.295 | 7536.145 | -3768.087 | -4.700 | -5.898 | -3.502 | 859.030 | 859.246 | -429.607 |
| M23: linear year x county + El Nino binary | 7538.720 | 51.336 | 7537.275 | -3768.659 | -4.956 | -6.247 | -3.666 | 861.134 | 860.376 | -430.180 |
| M5: linear year x county slope | 7544.185 | 50.106 | 7542.759 | -3771.400 | -4.700 | -5.978 | -3.422 | 866.599 | 865.860 | -432.921 |

**Table S3. Comparison of Bayesian spatiotemporal INLA models predicting the last day of the WNV transmission season in NYS.** Models differed in their temporal trend specification (linear or quadratic year), spatial covariates (elevation, latitude, or county-specific effects), and climate covariates (no ENSO term, linear ENSO effect, binary El Niño indicator, or a second-order random walk (RW2) representation of ENSO). Model performance was evaluated using the Deviance Information Criterion (DIC), effective number of parameters (pD), Widely Applicable Information Criterion (WAIC), and the log Conditional Predictive Ordinate (LCPO). Lower DIC and WAIC values indicate better model fit, while higher (less negative) LCPO values indicate better predictive performance. The estimated statewide change in the last day of the transmission season from 1999 to 2024 is reported as the posterior median (total_est) with 95% credible intervals (total_lo–total_hi). Positive estimates indicate a later end to the transmission season, whereas negative estimates indicate an earlier seasonal end. ΔDIC, ΔWAIC, and ΔLCPO represent differences relative to the best-performing model (M12).

| **model** | **DIC** | **pD** | **WAIC** | **LCPO** | **total_est** | **total_lo** | **total_hi** | **dDIC** | **dWAIC** | **dLCPO** |
| --- | --- | --- | --- | --- | --- | --- | --- | --- | --- | --- |
| M12: quadratic year x elev + ENSO RW2 | 5822.892 | 60.751 | 5824.832 | -2912.451 | 14.974 | 12.752 | 17.194 | 0.000 | 0.000 | 0.000 |
| M16: quadratic year x lat + ENSO RW2 | 5856.856 | 72.571 | 5859.102 | -2929.604 | 14.974 | 12.743 | 17.203 | 33.963 | 34.270 | -17.153 |
| M4: quadratic year + ENSO RW2 | 5858.389 | 70.270 | 5859.868 | -2929.981 | 14.982 | 12.751 | 17.212 | 35.497 | 35.036 | -17.530 |
| M8: quadratic year + county + ENSO RW2 | 5862.910 | 85.536 | 5866.794 | -2933.509 | 14.970 | 12.667 | 17.270 | 40.018 | 41.963 | -21.058 |
| M19: linear year x elevation + ENSO RW2 | 5916.741 | 58.083 | 5918.209 | -2959.135 | 7.338 | 6.384 | 8.290 | 93.849 | 93.378 | -46.684 |
| M20: linear year x latitude + ENSO RW2 | 5942.714 | 70.382 | 5944.279 | -2972.187 | 7.344 | 6.388 | 8.297 | 119.822 | 119.447 | -59.736 |
| M17: linear year + ENSO RW2 | 5944.906 | 69.435 | 5946.115 | -2973.102 | 7.344 | 6.388 | 8.298 | 122.014 | 121.283 | -60.651 |
| M18: linear year x county + ENSO RW2 | 5949.216 | 83.025 | 5953.260 | -2976.730 | 7.342 | 6.321 | 8.360 | 126.324 | 128.428 | -64.279 |
| M10: quadratic year x elevation | 6309.300 | 40.137 | 6310.466 | -3155.245 | 17.970 | 16.073 | 19.866 | 486.408 | 485.634 | -242.794 |
| M26: quadratic year x elevation + El Nino binary | 6312.403 | 41.911 | 6313.437 | -3156.732 | 17.969 | 16.070 | 19.868 | 489.511 | 488.605 | -244.281 |
| M14: quadratic year x latitude | 6344.031 | 53.316 | 6345.443 | -3172.744 | 17.970 | 16.068 | 19.871 | 521.139 | 520.611 | -260.293 |
| M2: quadratic year | 6344.275 | 50.985 | 6345.082 | -3172.560 | 17.970 | 16.064 | 19.875 | 521.382 | 520.250 | -260.109 |
| M28: quadratic year x latitude + El Nino binary | 6345.864 | 54.290 | 6347.306 | -3173.677 | 17.969 | 16.066 | 19.872 | 522.971 | 522.474 | -261.226 |
| M22: quadratic year + El Nino binary | 6346.134 | 51.971 | 6346.981 | -3173.511 | 17.969 | 16.062 | 19.876 | 523.241 | 522.150 | -261.060 |
| M6: quadratic year + county slope | 6351.225 | 63.268 | 6353.380 | -3176.740 | 17.970 | 15.980 | 19.959 | 528.332 | 528.548 | -264.288 |
| M24: quadratic year + county slope + El Nino binary | 6352.695 | 63.821 | 6355.325 | -3177.714 | 17.969 | 15.977 | 19.961 | 529.803 | 530.493 | -265.263 |
| M9: linear year x elevation | 6513.867 | 37.852 | 6514.559 | -3257.290 | 9.366 | 8.619 | 10.113 | 690.974 | 689.728 | -344.839 |
| M25: linear year x elevation + El Nino binary | 6515.218 | 38.823 | 6515.986 | -3258.004 | 9.379 | 8.632 | 10.126 | 692.325 | 691.154 | -345.553 |
| M11: linear year x elevation + ENSO linear | 6515.946 | 38.905 | 6516.511 | -3258.267 | 9.367 | 8.619 | 10.115 | 693.053 | 691.679 | -345.816 |
| M13: linear year x latitude | 6540.743 | 50.481 | 6541.699 | -3270.868 | 9.366 | 8.618 | 10.115 | 717.851 | 716.867 | -358.417 |
| M1: linear year | 6541.590 | 49.197 | 6542.280 | -3271.157 | 9.366 | 8.617 | 10.115 | 718.698 | 717.448 | -358.706 |
| M27: linear year x latitude + El Nino binary | 6542.031 | 51.480 | 6543.080 | -3271.560 | 9.379 | 8.631 | 10.128 | 719.139 | 718.248 | -359.109 |
| M15: linear year x latitude + ENSO linear | 6542.822 | 51.664 | 6543.667 | -3271.854 | 9.367 | 8.617 | 10.117 | 719.929 | 718.836 | -359.403 |
| M21: linear year + El Nino binary | 6543.026 | 50.193 | 6543.791 | -3271.914 | 9.379 | 8.630 | 10.129 | 720.133 | 718.959 | -359.463 |
| M3: linear year + ENSO linear | 6543.565 | 50.157 | 6544.141 | -3272.088 | 9.367 | 8.617 | 10.117 | 720.672 | 719.309 | -359.637 |
| M5: linear year x county slope | 6548.549 | 60.054 | 6551.489 | -3275.791 | 9.366 | 8.538 | 10.194 | 725.656 | 726.657 | -363.340 |
| M23: linear year x county slope + El Nino binary | 6550.799 | 61.956 | 6553.463 | -3276.781 | 9.379 | 8.545 | 10.213 | 727.906 | 728.631 | -364.330 |
| M7: linear year x county + ENSO linear | 6551.046 | 61.510 | 6553.447 | -3276.771 | 9.367 | 8.536 | 10.198 | 728.154 | 728.615 | -364.320 |

**Table S4. Paired comparisons of observed and climate-model–estimated season metrics. Wilcoxon signed-rank tests were used to compare paired GCM estimates with observed values for season length, first day, and last day**. n represents the number of paired GCM comparisons; median difference represents the median difference between paired estimates and observations. V and p values correspond to the Wilcoxon signed-rank test.

| **metric** | **n_gcm_pairs** | **median_diff** | **V** | **p_value** |
| --- | --- | --- | --- | --- |
| season_length | 37 | 26.7853346 | 701 | 1.40E-07 |
| first_day | 35 | -15.225474 | 15 | 9.32E-07 |
| last_day | 37 | 10.3888186 | 688 | 4.00E-07 |

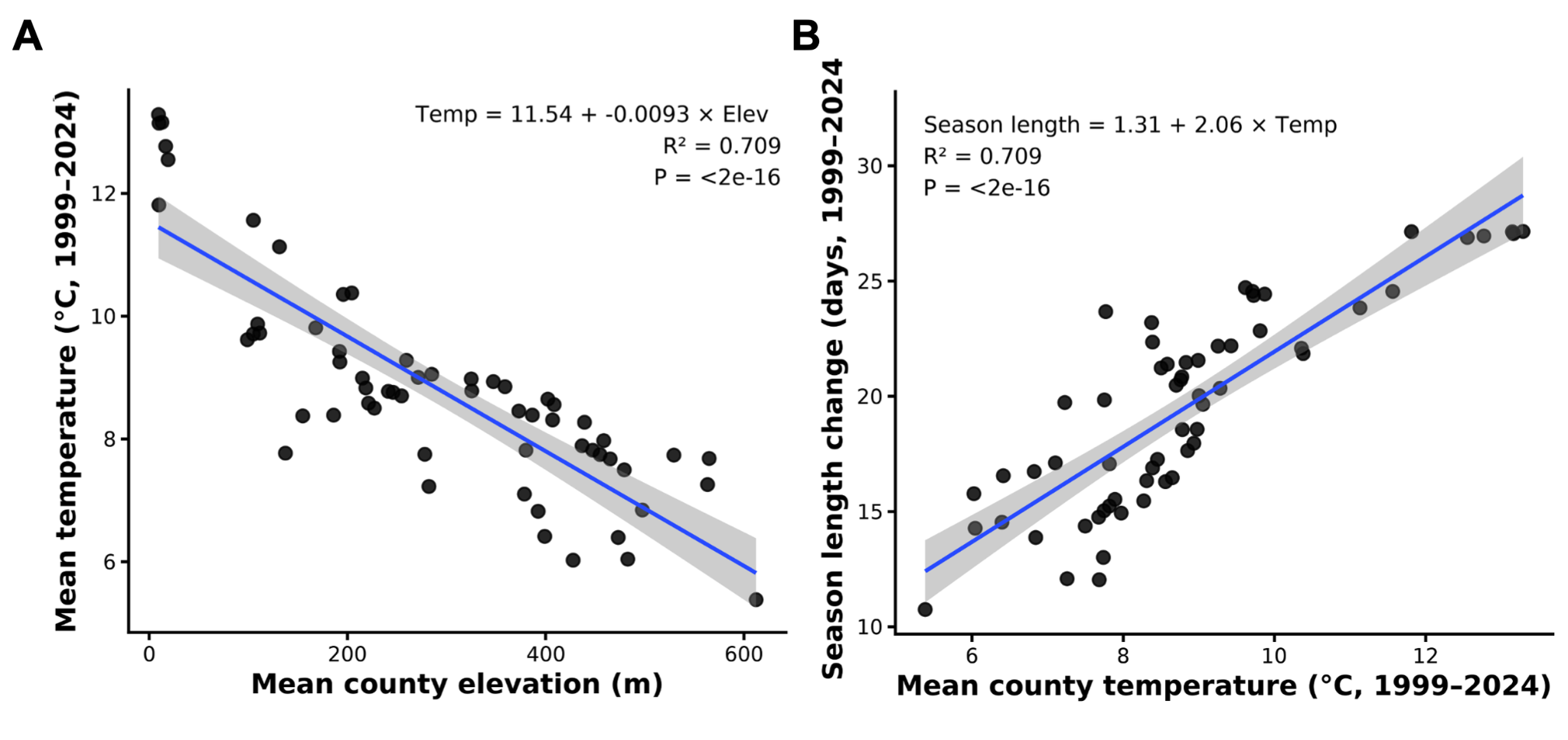

**Figure S1. Relationship between elevation and average change in county season length. A)** Mean county elevation and mean temperature from 1999-2024 have a significant relationship, 95% confidence interval shown in shading (simple linear regression, p < 0.0001). **B**) Relationship between elevation and change in WNV transmission season length (1999–2024) (linear regression model, p < 0.0001).

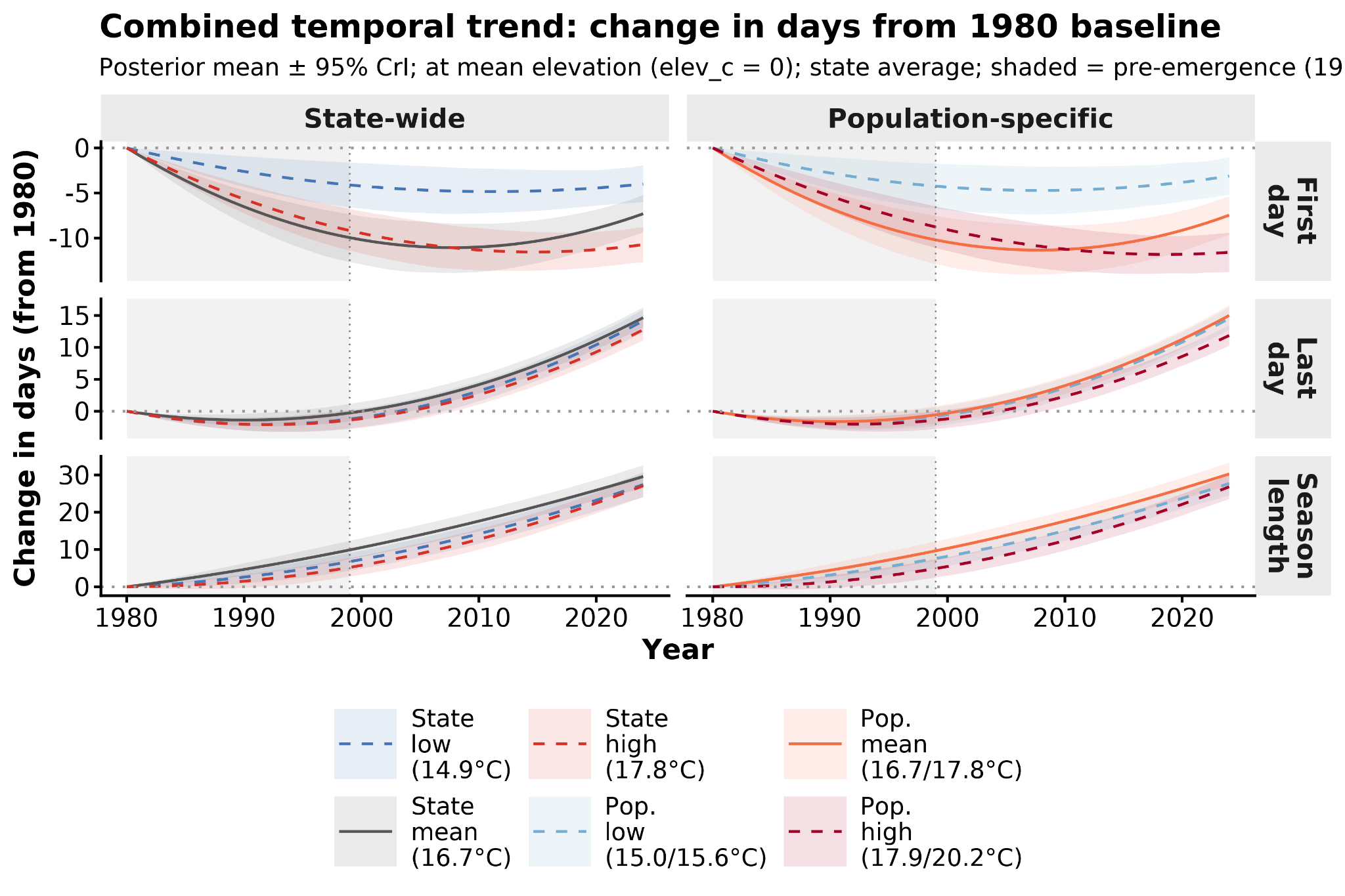

**Figure S2. Sensitivity of model results to temperature threshold. Rows show the first day of season onset (top), last day of season end (middle), and total season length (bottom).** The left column shows results using state-wide thermal thresholds; the right column shows results using population-specific thresholds. Lines show posterior means ± 95% credible intervals from final INLA models evaluated at mean elevation (elev_c = 0). Colors indicate the thermal suitability threshold used to define the transmission season: state-wide mean (16.7°C, black), state-wide low (14.9°C, blue dashed), state-wide high (17.8°C, red dashed), population-specific mean (16.7/17.8°C, orange), population-specific low (15.0/15.6°C, light blue dashed), and population-specific high (17.9/20.2°C, dark red dashed). Grey shading indicates the pre-WNV-emergence period (1980–1998). Consistency of trends across thermal thresholds indicates robustness of results to the choice of temperature cutoff.

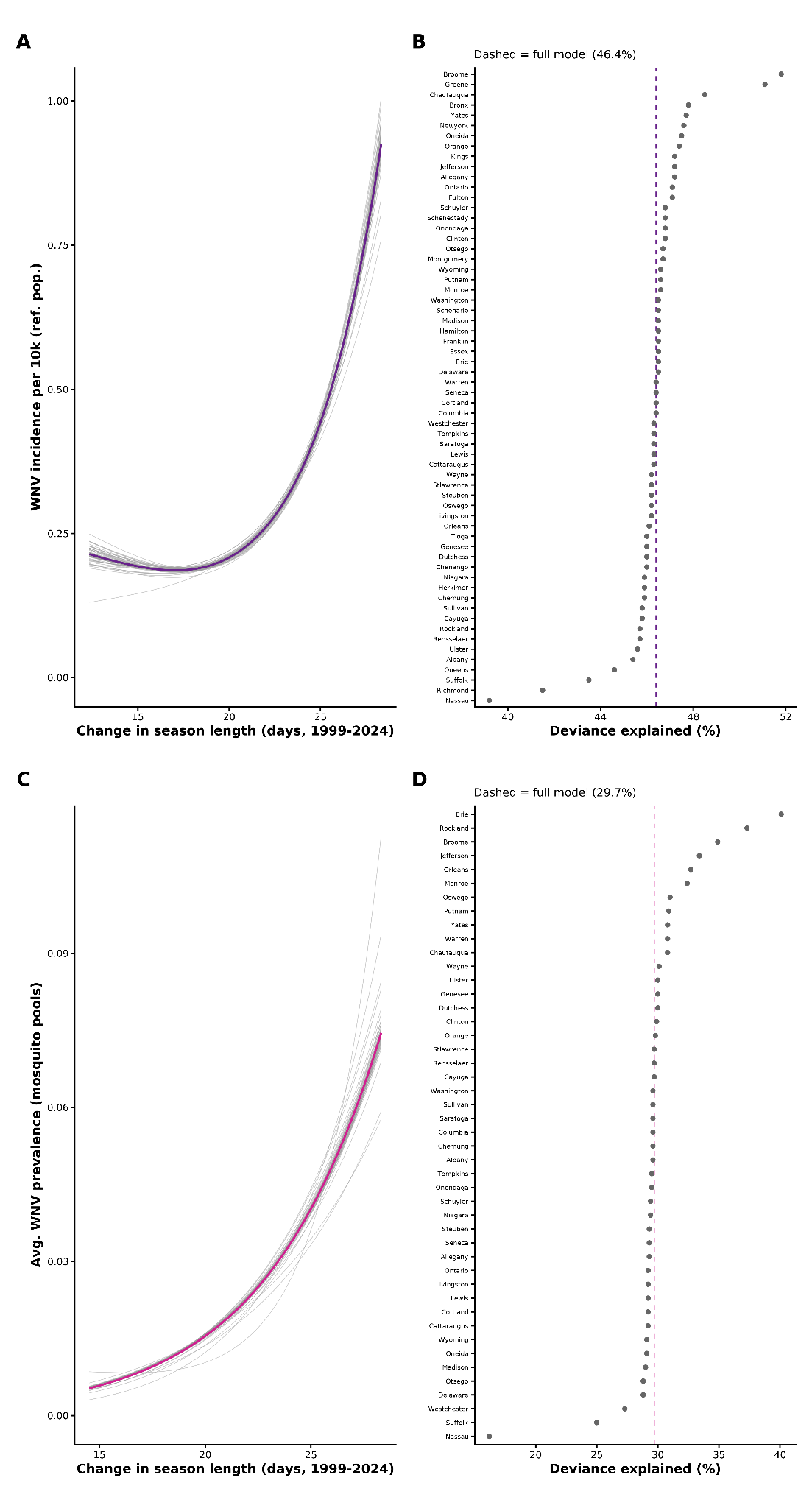

**Figure S3.** **Leave-one-out (LOO) sensitivity analysis for generalized additive models of WNV incidence and mosquito pool prevalence.** **A**) LOO smooth curves for the WNV human case incidence model (negative binomial with log-population offset); each grey line shows the GAM fit with one county excluded, and the purple line shows the full model fit. **B**) Deviance explained by the season length smooth for each LOO iteration of the incidence model, ordered from least to greatest; dashed purple line indicates full model deviance explained (46.4%). **C**) LOO smooth curves for the WNV mosquito pool prevalence model (quasibinomial); grey lines show fits with each county excluded, and the dark red line shows the full model. **D**) Deviance explained for each LOO iteration of the prevalence model; dashed red line indicates full model deviance explained (29.7%).

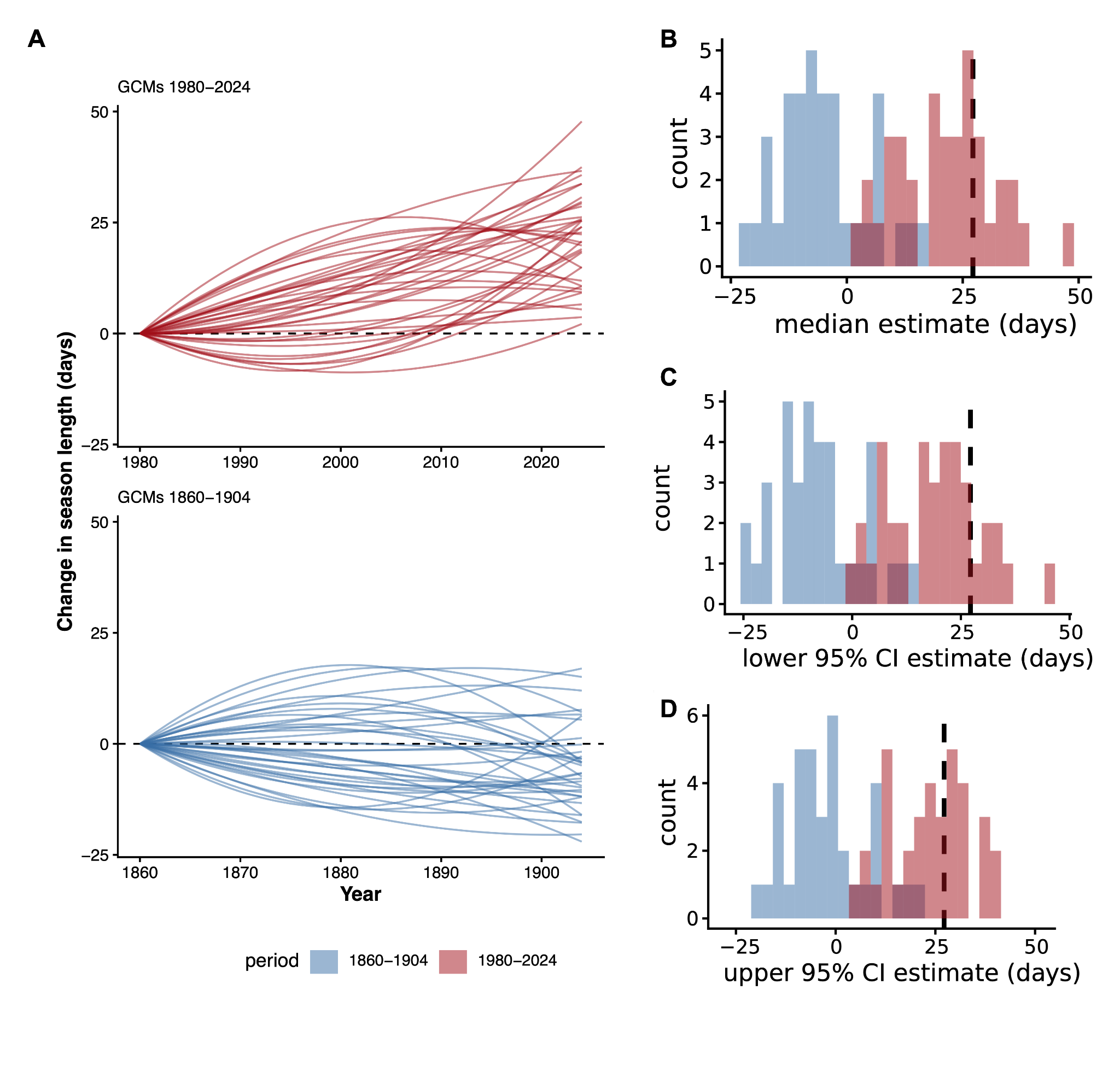

**Figure S4. Modeled change in WNV transmission season length under the 1980-2024 (modern) and 1860-1904 (pre-industrial-era analog) periods, using an ensemble of 37 downscaled GCMs**. **(A)** Predicted trajectories of change in season length (days), relative to the first year of each period, for each GCM individually, evaluated at mean county elevation (top: 1980-2024; bottom: 1860-1904). **(B-D)** Histograms of each GCM's Median (**B**), lower 95% CI (**C**), and upper 95% CI (**D**) estimate of the 45-year change in season length, by period; the black dashed line in each panel marks the corresponding observed GridMET value.

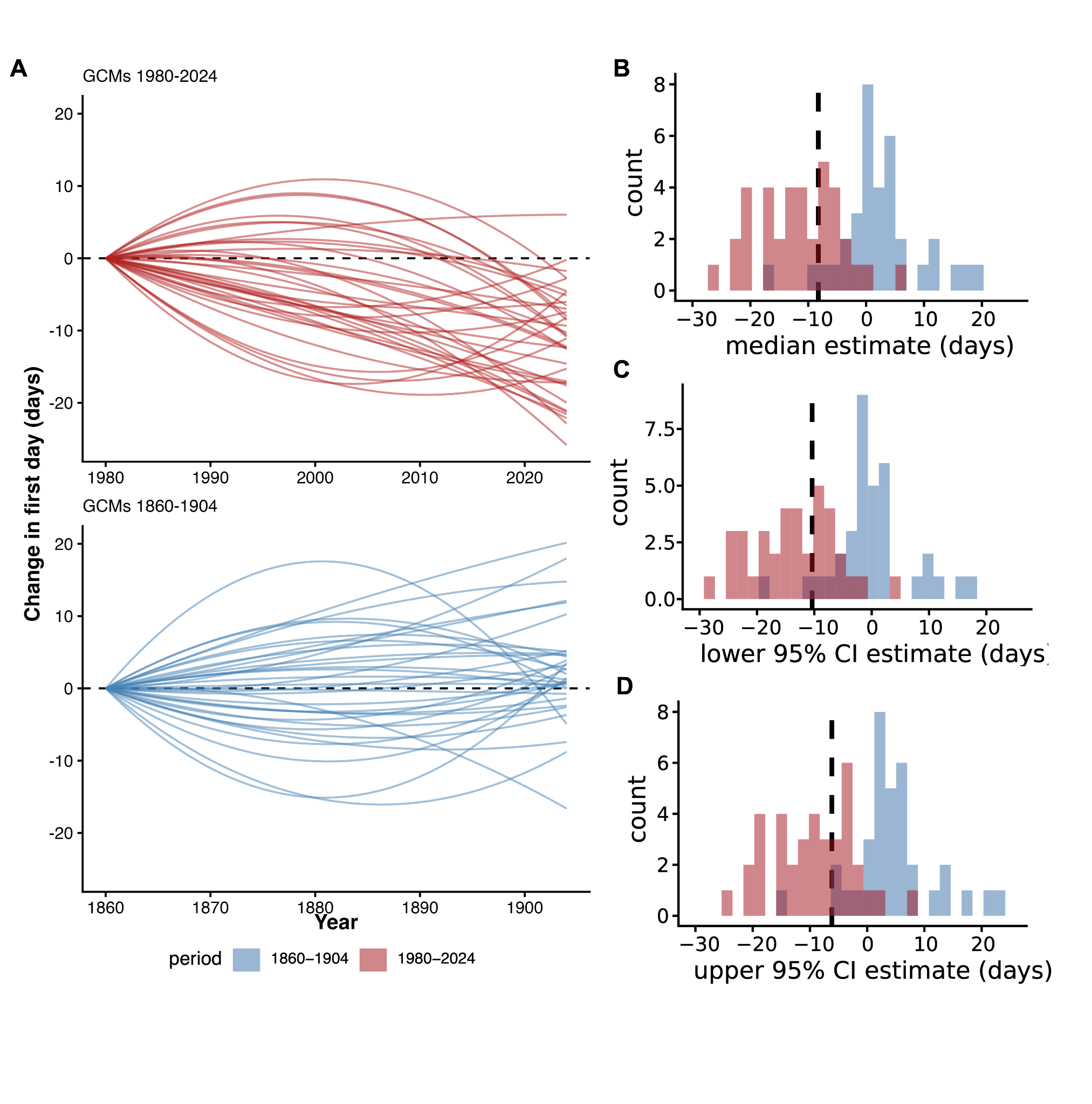

**Figure S5. Modeled change in WNV transmission start date under the 1980-2024 (modern) and 1860-1904 (pre-industrial-era analog) periods**, where a negative value indicates an earlier onset**. (A)** Per-GCM predicted trajectories of change in onset date (top: 1980-2024; bottom: 1860-1904). **(B-D)** Histograms of each GCM's Median (**B**), lower 95% CI (**C**), and upper 95% CI (**D**) estimate of the 45-year change in start date, by period; the black dashed line in each panel marks the corresponding observed GridMET value.

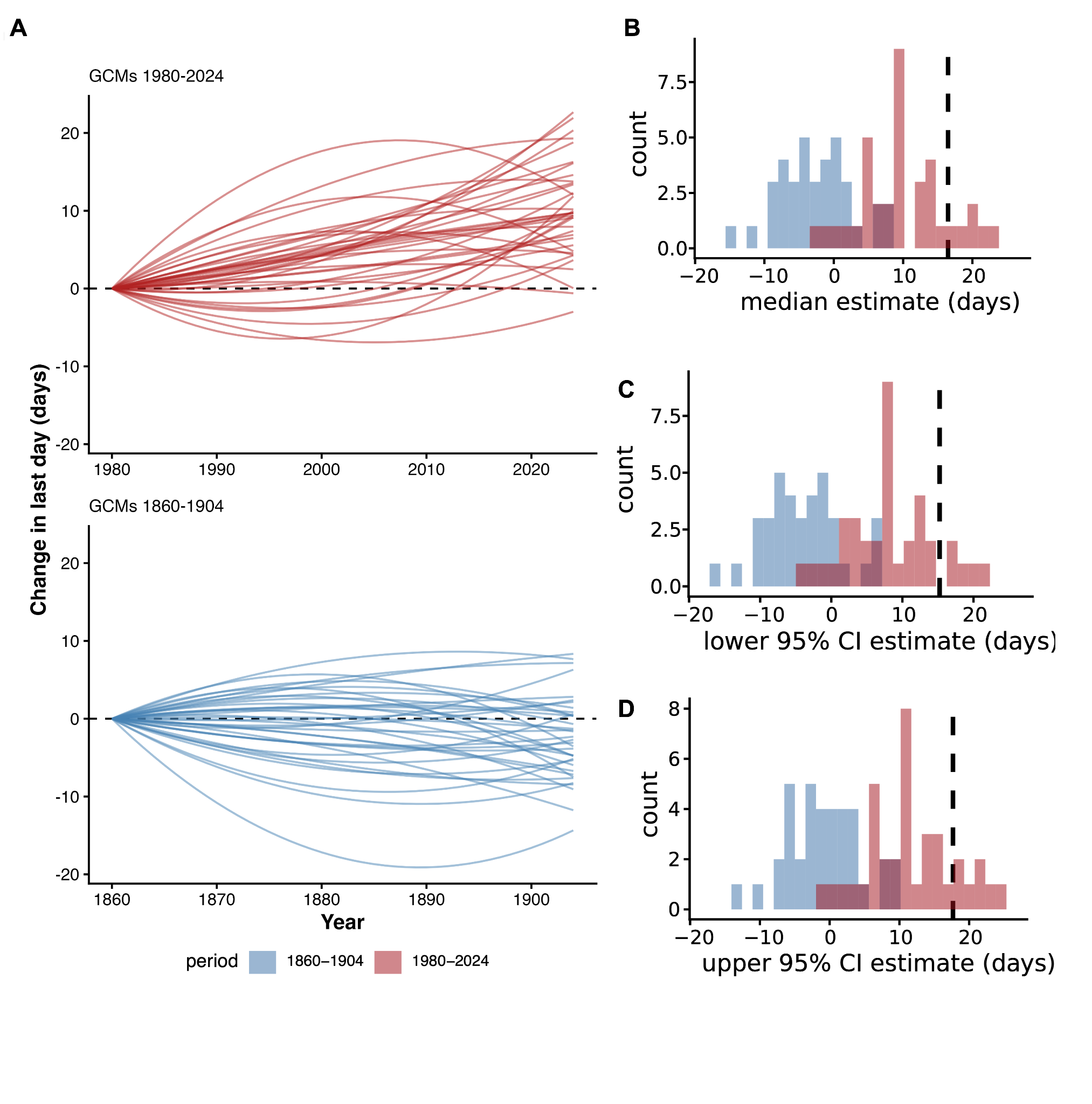

**Figure S6. Modeled change in WNV transmission end data date under the 1980-2024 (modern) and 1860-1904 (pre-industrial-era analog) periods**, where a positive value indicates a later last day of temperature suitable transmission. **(A)** Per-GCM predicted trajectories of change in termination date (Top: 1980-2024; Bottom: 1860-1904). **(B-D)** Histograms of each GCM's Median (**B**), lower 95% CI (**C**), and upper 95% CI (**D**) estimate of the 45-year change in termination date, by period; the black dashed line in each panel marks the corresponding observed GridMET value.

**Table S5.** Number and percentage of CMIP6 simulations with 45-year trends at least as large as the observed GridMET trend, for preindustrial (1860–1904) and modern (1980–2024) periods. Confidence intervals are one-sided 95% Clopper-Pearson upper bounds. Ratios compare modern to preindustrial probability. Where no preindustrial simulations met the observed trend, the ratio uses the one-sided upper bound (7.8% for 0/37) as the denominator and is a lower bound.

| **Metric** | **Period** | **N ≥ GridMET** | **% (95% CI, one-sided)** | **Ratio** | **N ≥ 0** | **% (95% CI, one-sided)** |
| --- | --- | --- | --- | --- | --- | --- |
| Season length | 1860-1904 (preindustrial) | 0 | 0.0% (≤7.8%) | 55.4/7.8 = 7.1x | 9 | 24.3% (≤38.6%) |
| Season length | 1980-2024 (modern) | 15 | 40.5% (≤55.4%) |  | 37 | 100% (≤100%) |
| First day (earlier) | 1860-1904 (preindustrial) | 2 | 5.4% (≤16.1%) | 75.5/16.1 = 4.7x | 9 | 24.3% (≤38.6%) |
| First day (earlier) | 1980-2024 (modern) | 23 | 62.2% (≤75.5%) |  | 36 | 97.3% (≤99.9%) |
| Last day | 1860-1904 (preindustrial) | 0 | 0.0% (≤7.8%) | 26.3/7.8 = 3.4x | 13 | 35.1% (≤50.0%) |
| Last day | 1980-2024 (modern) | 5 | 13.5% (≤26.3%) |  | 35 | 94.6% (≤99.0%) |
